# An interactive dashboard for global reports of the *Ralstonia solanacearum* species complex

**DOI:** 10.1101/2025.08.28.672672

**Authors:** Vienna N. Elmgreen, Benjamin Ramirez, Rituraj Sharma, Boris A. Vinatzer, Tiffany M. Lowe-Power, N. Tessa Pierce-Ward

## Abstract

The *Ralstonia solanacearum* species complex (RSSC) is a globally distributed group of gram-negative, soil-borne bacteria that cause wilt diseases on a broad range of hosts. Due to these pathogens’ impact on economically important plant species, there is a need for consolidated and visualized information on RSSC pathogen isolation data. We developed an interactive dashboard designed to allow users to explore and analyze the genetic diversity of the RSSC. The dashboard visualizes data in the form of maps, charts, and tables with a variety of user-interactive filters for taxonomic, geographic, and host of isolation specifications. This *Ralstonia* Wilt Dashboard will aid in communicating knowledge to researchers, regulatory scientists, and other stakeholders to improve disease control and regulation. This report highlights the deployment of the *Ralstonia* Wilt Dashboard and provides four case studies that address focused scientific questions (https://ralstoniadashboard.shinyapps.io/RalstoniaWiltDashboard/).

## Introduction

The *Ralstonia solanacearum* species complex (RSSC) is a globally distributed, diverse group of soil- and water-borne pathogens that cause wilt and rot diseases of many economically important plant species. RSSC are generalist pathogens, with individual strains having complex host range patterns^1,2^. Often, the RSSC pathogens that infect economically important crops can also cause latent infections of ornamental plants. Through movement of live plants, RSSC lineages have the potential and demonstrated ability to enter, establish, and spread to new locations^3,4^. Access to pathogen isolation data is vital for understanding pathogen distribution and improving disease management strategies to safeguard against impactful crop pathogens, but RSSC pathogen metadata remains difficult to access as the information is fragmented across the scientific literature. To investigate large-scale patterns in RSSC biogeography and host incidence, we previously developed a centralized resource of consolidated pathogen metadata, the RSSC Diversity Dataset / Database, a living pre-print and supplemental Excel spreadsheet with a stable digital object identifier (DOI) and version control^5^. Here we present the *Ralstonia* Wilt Dashboard, an interactive web-based dashboard that enables visualization of the full or filtered subsets of the RSSC Diversity Dataset / Database.

## Results and Discussion

The *Ralstonia* Wilt Dashboard has a simple interface with a left-hand sidebar and a central dashboard body (Fig. 1). The sidebar displays a brief description of the dashboard, the drop-down selections for filtering the dataset (Fig. 1A-B), buttons to apply or reset filters (Fig. 1C), and buttons to download the filtered or full datasets (Fig. 1D). At the top of the dashboard body, a display box directs users to provide feedback and report bugs via a Google Form (Fig. 1E). Four reactive information boxes display information based on selected filters: the number of isolates, research articles, countries, and hosts of isolation (Fig. 1F). A focal point of the *Ralstonia* Wilt Dashboard is a world map (Fig. 1G). This map displays the reported isolation locations of *Ralstonia*, and each data point is color-coded according to the isolate’s phylotype. The map has a toolbar with options to zoom in / out, pan around the map, download an image of the map, and reset the map to the default view (Fig. 1H). When a cursor hovers over a point on the map, a textbox displaying a brief overview of strain information appears (Fig. 1I). The map’s default jitter view can be toggled on or off with buttons below the map (Fig. 1J).

**Figure 1.**
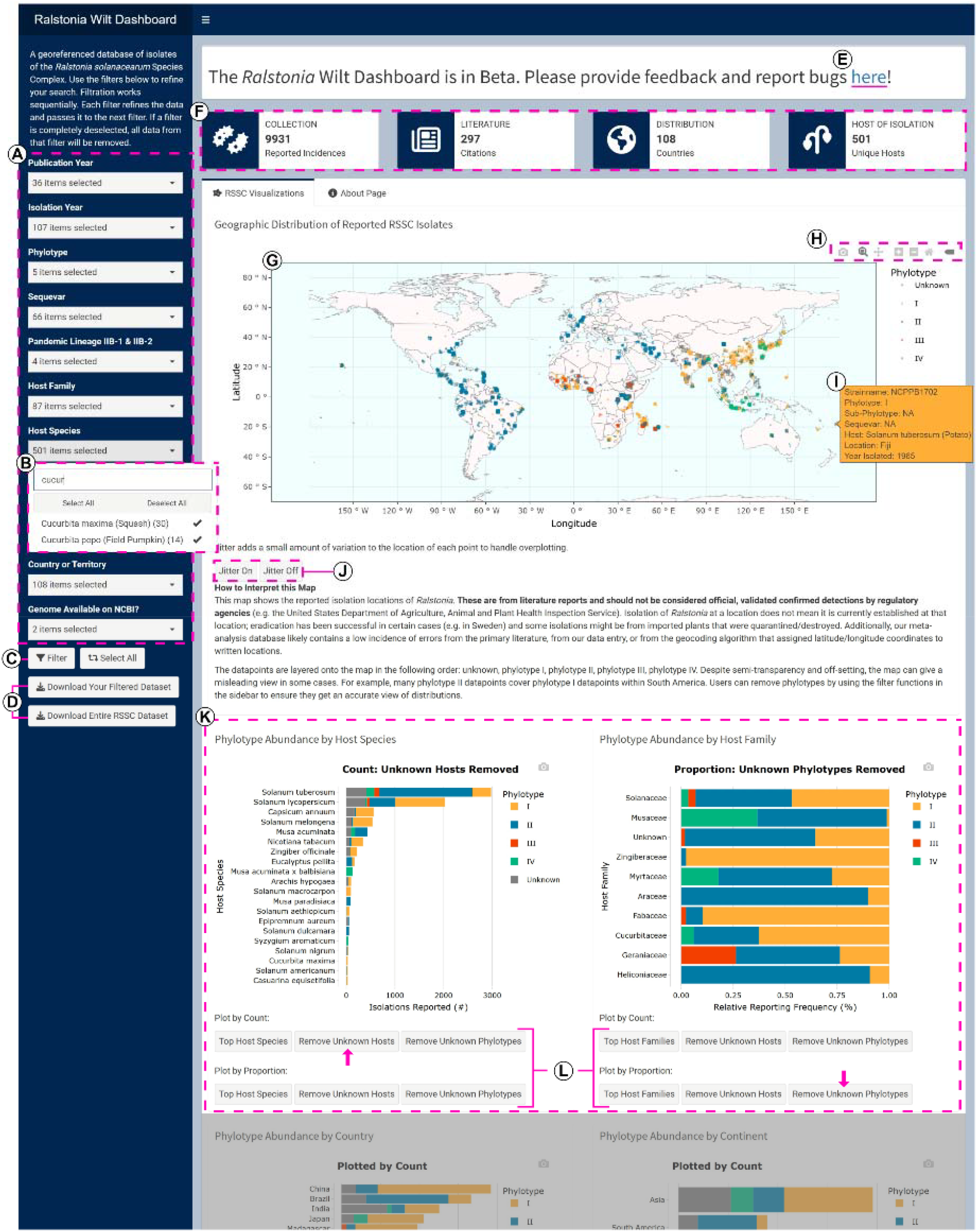
*Ralstonia* wilt dashboard user interface. Highlighted features: A) filter selections; B) dropdown search tool displaying an example query; C) button to apply filtered selections; D) buttons to download datasets in.csv format; E) link to the dashboard’s feedback and bugs form; F) information boxes that summarize the filtered dataset; G) map displaying the reported geographic distribution of isolates; H) interactives toolbar for the map; I) example of hover text displaying information about a reported strain; J) buttons to toggle the map’s jitter function on and off; K) charts displaying phylotype abundance by host of isolation; L) buttons to toggle between chart views, pink arrows indicate which buttons have been used to produce the chart. Some dashboard visualizations are not shown in this figure: two charts displaying phylotype abundance by location and a reactive data table displaying strain data.

Below the map, there are four charts that display phylotype abundance: two charts show host by species and by family (Fig. 1K) and two charts show location by country and by continent. Under each chart, there are buttons to toggle between views to exclude entries with unknown host or location metadata (Fig. 1L). Additionally, users can use these buttons to customize the charts to display phylotype abundance by the raw count or by proportion. At the bottom of the dashboard, there is a reactive data table that displays key information on filtered strains. The table has interactive search and sort functions, and users can download the full or filtered data as.csv by using the buttons on the lefthand sidebar (Fig. 1D).

The *Ralstonia* wilt dashboard visualizes a dataset of over 9,000 entries as of 2025. Despite data curation, errors in this large dataset may be present from the primary literature, from data entry, or from the geocoding pipeline used to assign latitude and longitude coordinates to text-based locations. Additionally, there are likely biases in the underlying data: RSSC reporting is impacted by funding availability and the prioritization of economically important crops. We hope to expand the *Ralstonia* Wilt Dashboard to include more visualizations and interactive features. Suggestions can be provided through the feedback form on the dashboard.

### Case studies: Using the dashboard to highlight biologically relevant patterns

The *Ralstonia* Wilt Dashboard can be used to explore strain distribution and diversity and observe patterns in hosts of isolation. The default view displays all entries in the dataset, which shows high-level patterns in phylotype distribution and host of isolation. The filters in the sidebar allow the user to tailor dashboard visuals with precision. Filtering with the dataset provides a closer look at RSSC distribution patterns, and surveillance of these patterns can shed light on the transmission of strains to new locations. Here we provide varied case studies that demonstrate how users can tailor the dashboard visuals to address focused scientific questions.

#### Case Study: Strains by Phylotypes

The RSSC phylotypes have unique geographic origins and vary in their present distributions. Phylotype I originated in Asia, phylotype II originated in the Americas, and currently both phylotype I and II are globally prevalent. Phylotype III originated in Africa, phylotype IV originated in Southeast Asia and Japan, and both phylotype III and IV are globally rare^6^. Fig. 2 shows four global maps that display the locations where strains of each phylotype have been isolated (Fig. 2A-D). These maps can be produced on the dashboard by filtering the dataset by each phylotype. Although RSSC isolations have been reported on over 500 plant species, the majority of reports are associated with potato and tomato (Fig. 2E). Phylotype I and phylotype II are most frequently reported in the literature, comprising 45% and 46% of reports in which phylotype is recorded, and these entries capture the largest breadth of plant species (Fig. 2F). Isolations associated with phylotype I and phylotype II have been reported on 126 and 82 unique hosts of isolation, respectively. Isolations associated with phylotype III and phylotype IV have only been reported on 16 and 20 unique hosts and comprise only 3% and 6% of reports, respectively.

**Figure 2.**
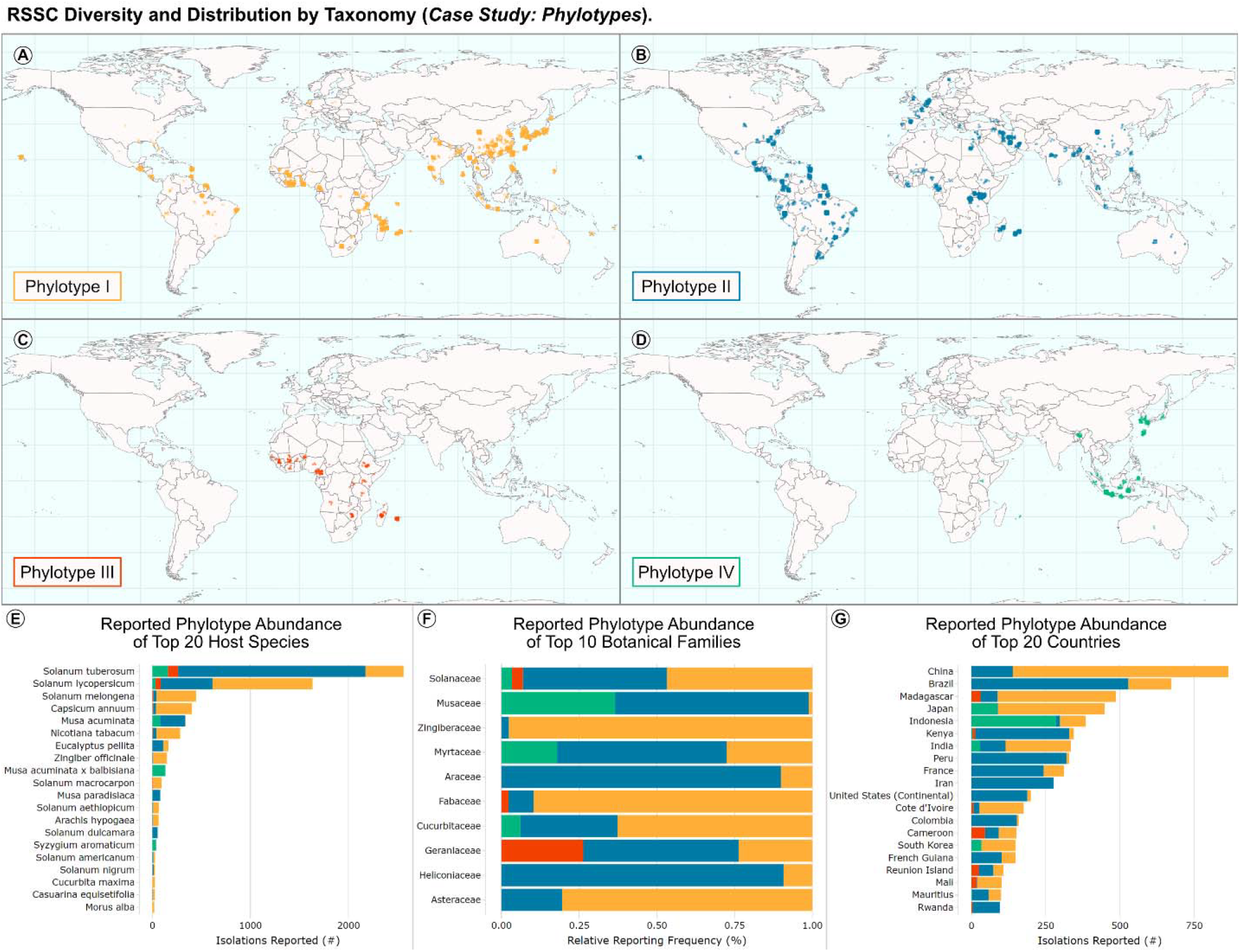
RSSC diversity and distribution (*Case Study: Phylotypes*). Maps display the reported strain distribution of A) phylotype I, B) phylotype II, C) phylotype III, and D) phylotype IV. The literature varied in reporting specificity for location (e.g. from city, province, country, etc.), so we employed Mapbox to infer latitude and longitude coordinates for the average center of the reported locations. The default view using jitter is shown; jitter randomly staggers symbols with identical geographical coordinates, yielding clusters that convey frequency of reports at specific locations. E-G) Stacked bar charts display the reported phylotype abundance of E) top 20 host species, F) top 10 botanical families, and G) top 20 countries. All chart data can be plotted either by count (E and G) or by proportion (F).

#### Case Study: Strains by Lineage (IIB-1 and IIB-2)

The dashboard can be used to investigate the distribution and reported hosts of individual RSSC lineages. In the United States, the closely related IIB-1 and IIB-2 lineages are regulated as a Select Agent under the historical name “race 3 biovar 2”. Dashboard visualizations can be tailored to highlight specific lineages (such as sequevars), including the IIB-1 and IIB-2 lineages (Fig. 3). IIB-1/IIB-2 strain isolations are reported from 61 countries (Fig. 3A) with the most abundant reporting from Kenya, Peru, and Iran, each with over 200 strains reported (Fig. 3D). Although isolations of IIB-1/IIB-2 strains have been reported on 23 plant species, potato (*Solanum tuberosum*), tomato (*Solanum lycopersicum*), and bittersweet nightshade (*Solanum dulcamara*) account for 90%, 4%, and 3% of the reported hosts of isolation (Fig. 3C). Users can similarly filter the data to explore patterns for other sequevars.

**Figure 3.**
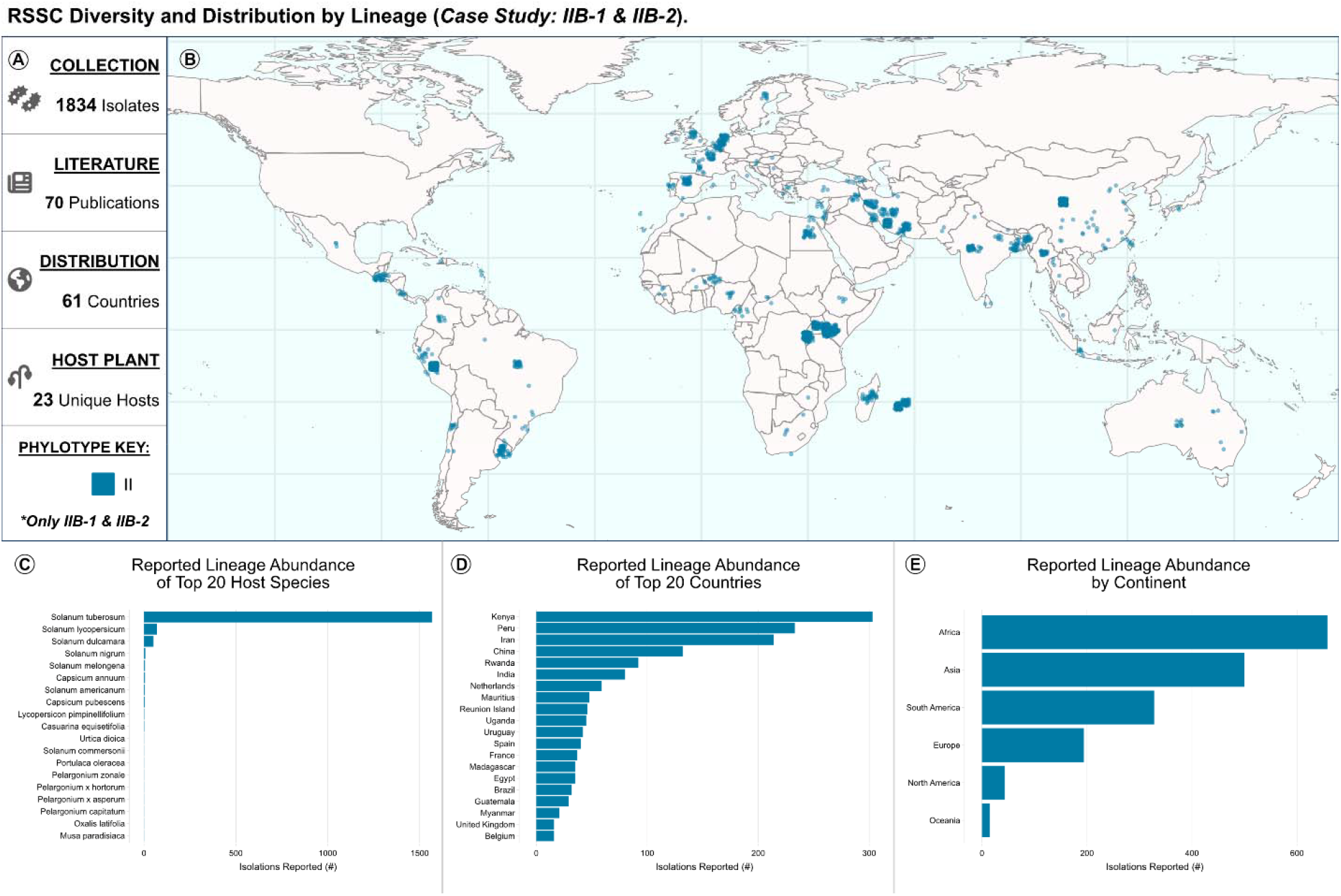
RSSC diversity and distribution by lineage (*Case Study: IIB-1* and *IIB-2*). A) Reactive information boxes summarize the filtered data: 1,834 isolates, 70 sources, 62 countries, and 24 unique host species. B) The map shows the reported geographic distribution of IIB-1 and IIB-2 isolates. C) IIB-1 and IIB-2 abundance by host species (count). D) IIB-1 and IIB-2 abundance by country (count). E) IIB-1 and IIB-2 abundance by continent (count).

#### Case Study: Strains by geographic location (Africa)

The dashboard visualizations can be tailored to focus on strains isolated from specific geographic locations, such as the African continent and nearby islands (Fig. 4). Although only phylotype III originated in Africa, all phylotypes are currently established in Africa. The map in Fig. 4B and chart in Fig. 4C demonstrate that phylotype I and II account for the majority of reported RSSC isolations in Africa. The two most thoroughly surveyed African countries are Madagascar and Kenya (Fig. 4E), with 487 and 351 reported strains. The next most thoroughly surveyed countries are Cote d’Ivoire, Ethiopia, Benin, and Cameroon, with over 150 reported strains from each country. RSSC pathogens have been isolated from 70 plant species in Africa (Fig. 4A), but potato (*Solanum tuberosum*) and tomato (*Solanum lycopersicum*) are most commonly reported, accounting for 40% and 33% of reports in which host of isolation is recorded. Users can similarly explore the patterns in RSSC reports from other continents or specific countries or territories.

**Figure 4.**
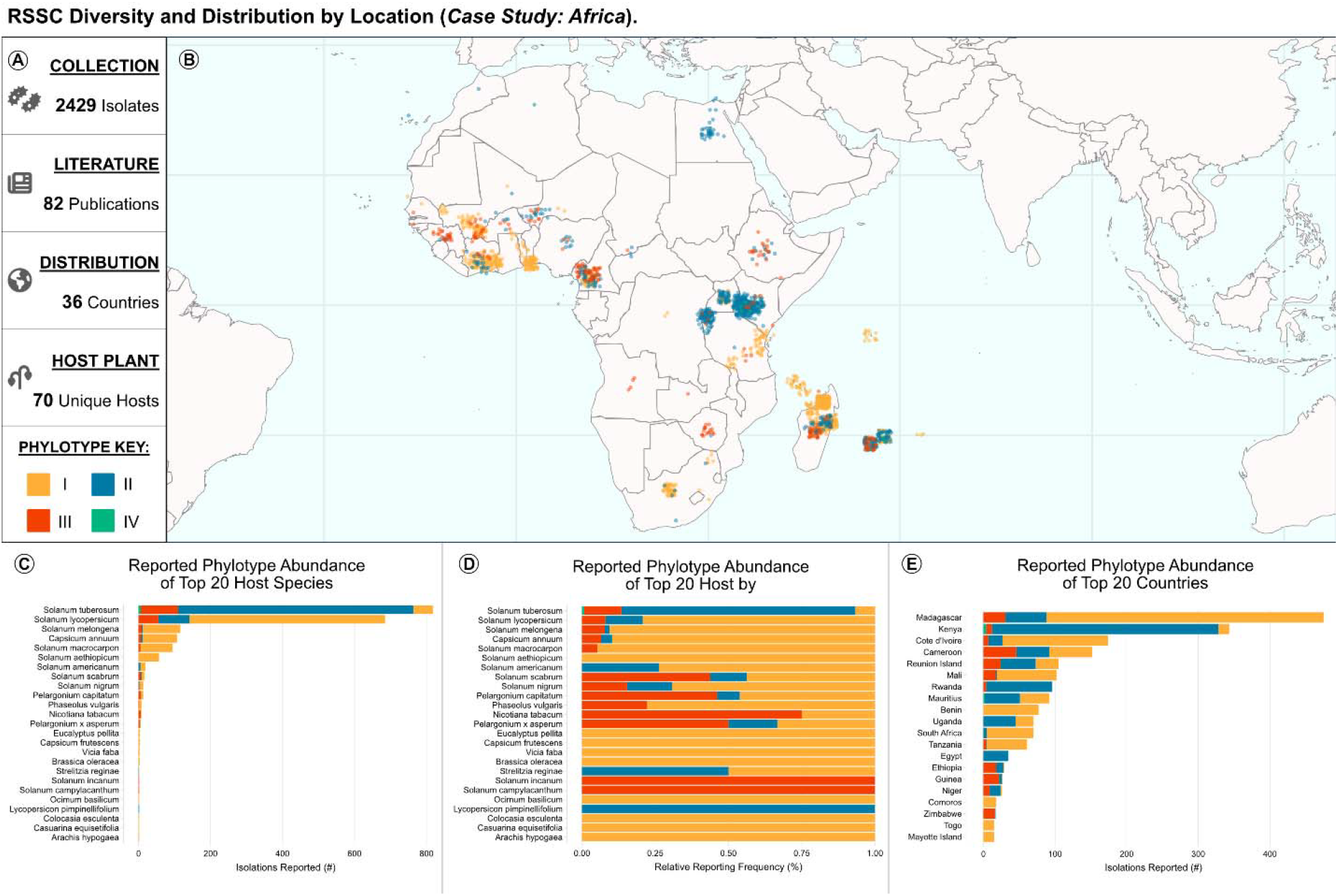
RSSC diversity and distribution by location (*Case Study: Africa*). A) Reactive information boxes summarize the filtered data: 2,449 isolates, 82 sources, 36 countries, and 70 unique host species. B) The map shows the reported geographic distribution of RSSC isolated in Africa. C) Phylotype abundance by host species (count). D) Phylotype abundance by host species (proportion). E) Phylotype abundance by country (count).

#### Case Study: Strains by host (Banana/plantain family)

The dashboard can be used to highlight patterns in reported host of isolation for RSSC strains. The data can be tailored to one-or-more specific plant families or plant species. Additionally, there is a drop-down filter specific to vegetatively propagated hosts such as potato, banana, and ginger. Strains isolated from Musaceae plants like banana and plantain are shown in Fig. 5. These data encompass 709 isolates from 61 sources (Fig. 5A). Since most banana cultivars are triploid seedless clones derived from hybridization of *Musa acuminata* and sometimes *Musa balbisiana*^7^, RSSC pathogens that have been reported on specific cultivars have been nested within “*Musa acuminata*” or “*Musa acuminata* x *balbisiana*”. The global map (Fig. 5B) and phylotype abundance by location charts (Fig. 5D,E) illustrate a well-documented distribution pattern^8–10^: phylotype II strains commonly infect *Musa* hosts in the Americas and phylotype IV strains commonly infect *Musa* hosts in Indonesia. Musaceae-associated RSSC strains have been reported in 26 countries (Fig. 5A) with the majority of strains reported from Indonesia, Colombia, and Brazil (Fig. 5D). Concerningly, the RSSC pathogens that cause banana blood disease (BBD) continue to expand geographically, with recent incidence reported in Thailand and Malaysia^11,12^.

**Figure 5.**
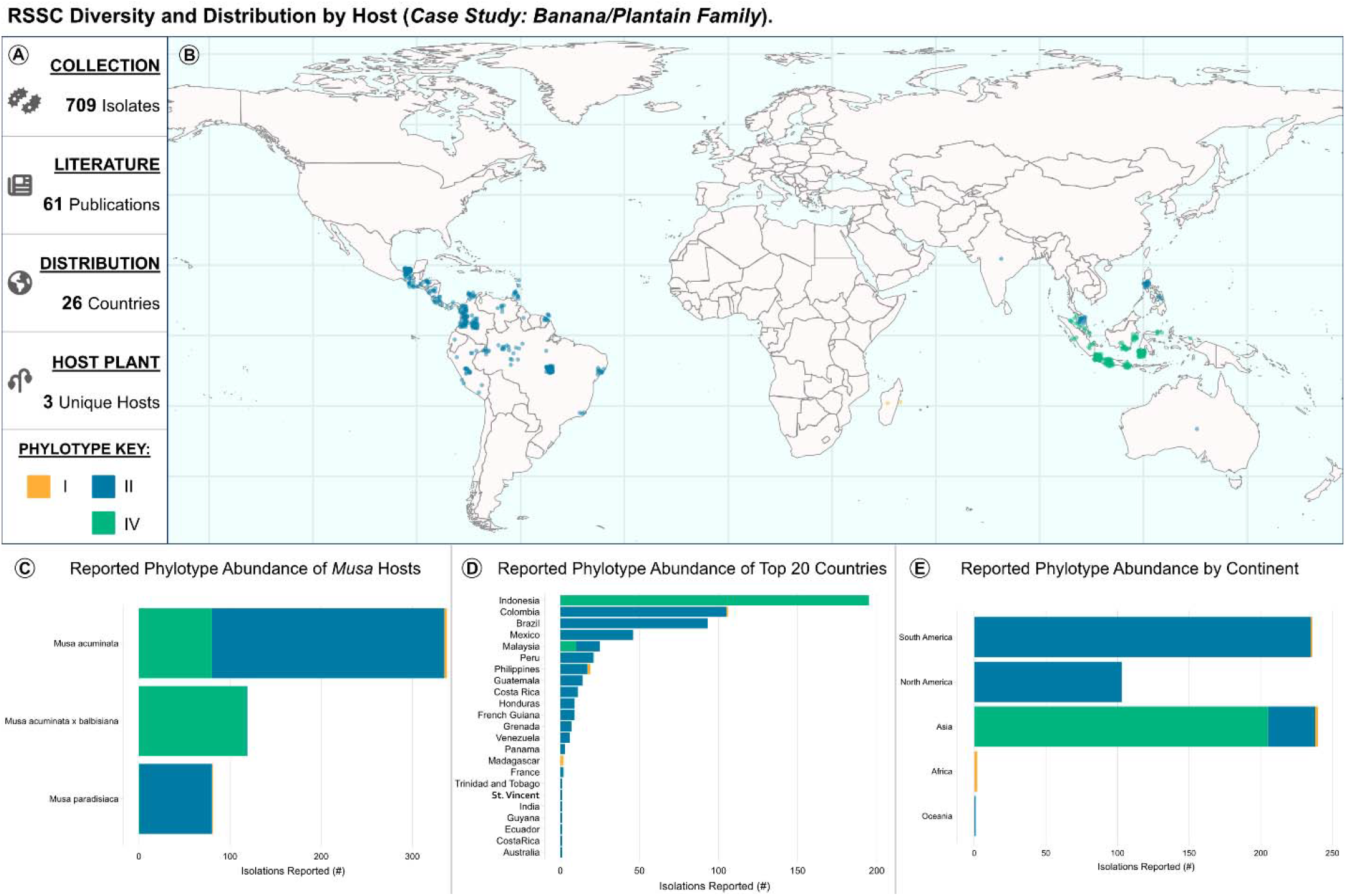
RSSC diversity and distribution by host (*Case Study: Banana/Plantain Family*). A) Reactive information boxes summarize the filtered data: 709 isolates, 61 sources, 26 countries, and 4 unique host species. B) The map shows the geographic distribution of reported entries isolated from Musaceae hosts. C) Phylotype abundance per *Musa* species (count). D) Phylotype abundance by country (count). E) Phylotype abundance by continent (count).

## Methods

We previously created the RSSC Diversity Dataset / Database, a centralized resource that consolidates and cites primary reports of the reported global distribution and host range of *Ralstonia* clades. The dataset and methods of data curation are described in detail in our living pre-print^5^. As of October 2024, the RSSC Diversity Dataset / Database catalogs information from nearly 10,000 strains from over 300 sources published from 1971 to 2024 that report one or more *Ralstonia* strains isolated from over 400 host plant species from over 100 geographic regions curated from over 300 sources.

### Data Curation

Data cleaning and standardization was performed manually in Excel and *en masse* with OpenRefine to ensure consistent naming and formatting across entries. OpenRefine is an open-source tool for working with messy datasets that allows users to cluster, clean, and transform data^13^. First, duplicate strain entries were manually identified and merged in Excel, then OpenRefine was used to normalize categorical entries of equivalent text. Longitude and latitude were assigned to all entries in which ‘Location Isolated’ data was available. A custom script with MapBox was used to determine approximate longitudinal and latitudinal coordinates of recorded locations; accuracy was confirmed by checking whether the computed coordinates fell within the correct associated continental bounding box. The RSSC database was copied to a Google Sheet that is version controlled and directly populates the *Ralstonia* Wilt Dashboard with the RSSC metadata.

### Data Summary and Development of the R Shiny App

We developed the *Ralstonia* Wilt Dashboard using an RShiny web application framework built in RStudio written in R, HTML, and CSS. The layout of this dashboard was inspired by the *Fusarium graminearum* species complex database^14^. Our app is structured as a single app.R script (RWdbApp.R) containing both the user interface (UI) and server logic. The layout uses a ‘dashboardPage()’ structure with filtering and download options displayed in the sidebar and data visualizations displayed in the body. The UI incorporates interactive inputs such as: ‘pickerInput()’ for filtering the dataset; ‘actionButton()’ for submitting and resetting filter selections; and ‘downloadButton()’ for downloading a filtered subset or the full dataset. The server relies on reactive expressions, ‘eventReactive()’ and ‘observeEvent()’, to calculate values and perform actions in response to user inputs. We used several R packages to create interactive, responsive visualizations: ggplot2 was used to generate a map with strain locations and stacked bar charts showing phylotype patterns; plotly was used to incorporate hover tooltips, zoom, and download options into visualizations; ‘DT()’ was used to display a reactive, searchable, and filterable data table; and several additional packages were used to control aesthetics and other minor functions.

The app is deployed with Shinyapps.io, a cloud platform for hosting R Shiny applications, where it runs in a protected environment with SSL-encrypted access. Deployment is facilitated using rsconnect. All data, R scripts, and code associated with the *Ralstonia* Wilt Dashboard is publicly available on the dashboard GitHub repository (https://github.com/lowepowerlab/dashboard) and version 1.0 has been released and assigned stable DOIs (code: doi.org/10.5281/zenodo.16945323 and dataset: doi.org/10.5281/zenodo.16973477). The *Ralstonia* Wilt Dashboard can be accessed at: https://ralstoniadashboard.shinyapps.io/RalstoniaWiltDashboard/ or run locally.

### Primary Functions

There are 11 filter options built into the sidebar: Publication Year, Isolation Year, Phylotype, Sequevar, Pandemic Lineages, Host Family, Host Species, Vegetatively Propagated (VP) Hosts, Continent, Country or Territory, and Genome Available on NCBI. Each filter has a dropdown selection with a search bar where users can select and deselect options for each filter. Filters are in “select all that apply” format with additional “select all” and “deselect all” buttons, allowing for precision when tailoring selections. Choices listed in the dropdown selection display the number of entries found in the dataset associated with that choice. The filters operate in a sequential manner corresponding to the order they are displayed in the sidebar. The first filter refines the dataset and passes it to the next filter, and so on, until the data have been passed through the final filter. Filters can only pass on data that have been selected. If any of the filter options are completely deselected, then none of the data are passed to the next filter, which results in blank visualizations. All filter options have been selected as the default. Two buttons are located below the last filter selections: (1) a “Filter” button that submits the selected filters, prompting reactive events to update the data visualizations to reflect the user’s specifications and (2) a “Select All” button that resets all filter selections. Two download buttons are located below these, a “Download Your Filtered Dataset” button that allows users download a.csv file containing the filtered RSSC metadata, and a “Download Entire Dataset” button that allows users to download a.csv file of the full RSSC Diversity Database dataset.

The dashboard body is structured using a ‘tabBox()’ with two tabs. The first tab, “RSSC Visualizations”, displays a map showing the geographic distribution of reported isolates, four stacked bar charts with multiple view options to display phylotype distribution patterns, and a metadata table. The second tab, “About Page”, displays an.Rmd file with additional information about the dashboard including purpose, how to use, how to report errors or bugs, about the team, acknowledgments, and links to related databases. This RSSC isolates map was built using ggplot and plotly with additional packages for layers and aesthetics which can be found in our source code. The map has several user-interactive features, such as zoom, pan, download, and hover text to display strain name, phylotype, sub-phylotype, sequevar, host, location, and year isolated. There are two buttons below the map that allow users to remove or apply jitter when plotting. Jitter adds a small amount of random variation to the placement of points when strains have identical coordinates. The purpose of this map is to display established RSSC strains isolated from field-grown plants, so we have endeavored to exclude data where RSSC strains were isolated from imported greenhouse plants.

Database entries with missing phylotype data are colored gray and labeled “Unknown”, and entries with missing ‘Location Isolated’ data have been removed from the map. We display the following disclaimer below the Geographic Distribution of Reported RSSC Isolates map:

*“This map shows the reported isolation locations of Ralstonia. These are from literature reports and should not be considered official, validated confirmed detections by regulatory agencies (e*.*g. the United States Department of Agriculture, Animal and Plant Health Inspection Service). Isolation of Ralstonia at a location does not mean it is currently established at that location; eradication has been successful in certain cases (e*.*g. in Sweden) and some isolations might be from imported plants that were quarantined/destroyed. Additionally, our meta-analysis database likely contains a low incidence of errors from the primary literature, from our data entry, or from the geocoding algorithm that assigned latitude/longitude coordinates to written locations*.*”*

In the RSSC Visualizations tab, under the map, there are four stacked bar charts that display patterns in phylotype abundance. The first two charts show host information at the species and family level, and the next two charts show location information at the country and continent level. Each visualization has an option to download it in vector (.svg) format. We applied the color-blind friendly palette “Egypt” from the MetBrewer R package to each visualization. Below the phylotype abundance charts, there is a reactive data table with built-in sorting and search functions. The data table displays an abridged version of the database, which includes index, phylotype, sequevar, strain name, host species (common name), host family, year isolated, location isolated, genome accession, and publication.

## Funding

Support was provided by the U.S. Department of Agriculture Hatch Projects (#1023861) and Pests and Beneficial Species in Agricultural Production Systems (A1112) program, project award no. 2024-67013-42781, from the U.S. Department of Agriculture’s National Institute of Food and Agriculture. Additionally, this material was made possible, in part, by PPA7721 Cooperative Agreements (AP24PPQS&T00C076 and AP25PPQS&T00C013) from the United States Department of Agriculture’s Animal and Plant Health Inspection Service (APHIS). It may not necessarily express APHIS’ views.

## Acknowledgements

We thank the global plant pathology community for investigating and reporting *Ralstonia* incidence and isolation information. We thank Emerson del Ponte (Federal University of Viçosa) for sharing advice and code from the FGSCdb which can be accessed at: https://edelponte.shinyapps.io/FGSCdb/.

## Author Contributions (CRediT)

VNE: Data curation, Analysis, Experiments / Investigation, Methodology, Software, Visualization, Writing – original draft, Writing – review & editing; BR: Data curation, Experiments / Investigation, Methodology, Software, Supervision, Writing – review & editing; RS: Experiments / Investigation, Methodology, Software, Visualization, Writing – review & editing; BAV: Funding acquisition, Methodology, Writing – review & editing; TMLP: Conceptualization, Data curation, Funding acquisition, Project administration, Supervision, Writing – review & editing; NTPW: Conceptualization, Data curation, Funding acquisition, Experiments / Investigation, Methodology, Project administration, Software, Supervision, Writing – review & editing

## Conflicts of Interest

The authors declare no conflicts of interest.

## Notes

https://ralstoniadashboard.shinyapps.io/RalstoniaWiltDashboard/

### Competing Interest Statement

The authors have declared no competing interest.

